# Multithreaded variant calling in elPrep 5

**DOI:** 10.1101/2020.12.11.421073

**Authors:** Charlotte Herzeel, Pascal Costanza, Dries Decap, Jan Fostier, Roel Wuyts, Wilfried Verachtert

**Author notes:** These authors contributed equally to this work (Charlotte Herzeel, Pascal Costanza).

## Abstract

We present elPrep 5, which updates the elPrep framework for processing sequencing alignment/map files with variant calling. elPrep 5 can now execute the full pipeline described by the GATK Best Practices for variant calling, which consists of PCR and optical duplicate marking, sorting by coordinate order, base quality score recalibration, and variant calling using the haplotype caller algorithm. elPrep 5 produces identical BAM and VCF output as GATK4 while significantly reducing the runtime by parallelizing and merging the execution of the pipeline steps. Our benchmarks show that elPrep 5 speeds up the runtime of the variant calling pipeline by a factor 8-16x on both whole-exome and whole-genome data while using the same hardware resources as GATK 4. This makes elPrep 5 a suitable drop-in replacement for GATK 4 when faster execution times are needed.

## Introduction

elPrep [1,2] is an established software tool for analyzing aligned sequencing data. It focuses on supporting community-defined standards such as the sequence alignment/map file format (SAM/BAM) [3] and the GATK Best Practice pipelines [4,5] for storing and analyzing sequencing data respectively. The main difference between elPrep and other tools for processing this kind of data such as Picard, SAMtools [3], and GATK 4 [6] lies in its software architecture that parallelizes and merges the execution of the pipeline steps while minimizing the number of data accesses to files. Our previous work [1,2,7] shows that this design greatly speeds up the runtimes of both whole-genome and whole-exome pipelines.

Previous versions of elPrep focus on preparation steps that prepare the data for statistical analysis by variant calling algorithms. This includes, among others, all of the preparation steps from the GATK Best Practice pipelines for variant calling [4], including steps for sorting reads, PCR and optical duplicate marking, and base quality score recalibration and application. The challenge to add new steps to elPrep is that the underlying algorithms need to be adapted so they fit into elPrep’s execution engine that parallelizes and merges computations. We additionally always guarantee that the output elPrep produces for any step is identical to the output of the reference tool, for example GATK 4, generates. This creates additional complexity from the implementation side, leading us to develop multiple new algorithms [1,2,7]. From a user’s perspective, however, it makes elPrep a drop-in replacement for other tools, resulting in its adoption by different bioinformatics projects [8–15].

The new elPrep 5 release introduces variant calling based on the haplotype caller algorithm from GATK 4 [6]. After optimizing the preparation steps, which allows elPrep to reduce the runtime per sample by many hours [2], variant calling remains a step that takes up a large chunk of the overall runtime of a pipeline. Looking at benchmarks, we see that, depending on the pipeline configurations and data inputs, variant calling takes up between 38% and 80% of the runtime when executing the pipeline with GATK 4. When executing the preparation steps with our previous version 4 release of elPrep, their runtime substantially reduces so that the variant calling step dominates and takes up between 82% and 93% of the total runtime. This suggests that adding variant calling allows to further increase the impact of elPrep on the overall runtime of a pipeline.

In the rest of this paper, we discuss the challenges and impact of integrating the haplotype caller algorithm into the elPrep framework. We analyze benchmarks that show we achieve similar speedups as to what we previously obtained for optimizing the preparation steps of the variant calling pipeline from the GATK Best Practices [1,2]. Concretely, we observe between 8-16x speedup compared to GATK 4 when using the same hardware resources.

## Materials and methods

We next describe the elPrep 5 software, the benchmark experiments, data sets, software versions, and hardware setups we use to validate our extensions to elPrep.

### Implementation

elPrep is developed for Linux at the ExaScience Lab at imec. The elPrep software is written in Go, a programming language developed by Google. The software is released with a dual license, both as an open-source project on GitHub (https://github.com/exascience/elprep) and the option for a premium license with support. There is otherwise (feature-wise or performance-wise) no difference between the two versions.

### elPrep 5 overview

The main new feature in elPrep 5 is the addition of variant calling based on the haplotype caller algorithm from GATK 4 [6]. We implement all three output modes, with GVCF as default, as well as various parameters to configure the variant calling process, which are all described in detail in the elPrep user manual.

#### Command-line interface with haplotype caller

The elPrep 5 software is distributed as a single executable for Linux. Formulating a pipeline in elPrep is achieved by means of a single command-line call. Listing 1 shows the elPrep invocation for implementing the variant calling pipeline from the GATK Best Practices.

elprep sfm NA12878.input.bam NA12878.output.bam

––mark – duplicates
––mark-optical – duplicates NA12878.output.metrics
––sorting – order coordinate
––bqsr NA12878.output.recal
––known – si tes dbsnp_138.hg38. elsites, Mills.hg38.elsites
––reference hg38.elfasta
–– haplotypecaller NA12878.output.vcf.gz **Listing 1.** elPrep command for the GATK Best Practices variant calling pipeline.

The command in Listing1 executes a pipeline that takes as input aligned sequencing data in the form of a BAM file (NA12878.input.bam), performs a number of data preparation steps, writes the output from those steps to a new BAM file (NA12878.output.bam), and finally performs variant calling, writing the output to a compressed VCF file (NA12878.output.vcf.gz). The preparation steps include PCR and optical duplicate marking with metrics logging, sorting by coordinate order, and base quality score recalibration taking into account known variant sites and the reference genome.

Additional parameters are possible for all of the options used in the command, but these are not listed here. The order in which the pipeline steps are listed is irrelevant. The elPrep implementation internally takes care of the correct execution order, while possibly merging and parallelizing the computations to speed up the execution. elPrep uses its own data formats for representing reference files (cf. elfasta) and files for storing known variant sites (cf. elsites). These are generated by separate elPrep commands once from the FASTA and VCF files that are normally used for such data. We opted for our own formats here because they can be parsed significantly more efficiently than plain VCF and FASTA data [2]. For more details, we refer to our extensive documentation online (http://github.com/ExaScience/elprep).

#### Framework extensions for haplotype caller

The haplotype caller is a non-trivial extension to the elPrep framework. To give an idea of the complexity, the haplotype caller code is roughly three times the code for the base quality score recalibration, previously the biggest extension to elPrep [2]. Adding variant calling also challenges the original architecture of the elPrep implementation.

The elPrep execution engine is a phased architecture with multiple intercession points where computations on the SAM/BAM data can be injected [1,2]. For example, there are different phases in the execution where filters that express operations on individual reads can be executed and phases where operations on the whole set of reads can be expressed. Concrete filters and operations are implemented separately from the engine using higher-order functions, cleanly separating the execution engine from the implementation of pipeline steps [1, 2]. elPrep is designed this way because it is intended to be an open-ended framework, encouraging the development of new filters and operations that intercept the execution engine.

For variant calling, we extended the phases in the elPrep execution engine with operations that execute on contigs of reads rather than individual reads or all of the reads at the same time. We also parallelized different parts of the 3-phase haplotype caller algorithm, which required restructuring the algorithms and adding proper synchronization points to avoid generating different results from GATK 4. For more details on the technical challenges we solved for adding variant calling, please consult the S1 Appendix.

### Benchmarks

To investigate the performance impact of adding variant calling, we benchmark the GATK Best Practice pipeline for variant calling [4, 5] on different data sets and hardware setups.

#### Pipeline

The concrete variant calling pipeline that we benchmark consists of the five steps listed below. As a reference, we list the GATK 4 tool name for each step between brackets:

1. Sorting the input BAM by coordinate order (SortSam);
2. Marking optical and PCR duplicates (MarkDuplicates);
3. Recalibrating base quality scores (BaseRecalibrator);
4. Applying base quality score recalibration (ApplyBQSR);
5. Variant calling using the haplotype caller algorithm (HaplotypeCaller).

#### Experiments

We set up two main types of benchmark experiments to investigate our computational performance claims. First, we present a detailed comparison of runtime, RAM, and disk use between elPrep 5 and GATK 4 when executing the variant calling pipeline. We subsequently present a scaling experiment on Amazon Web Services where we compare the parallel scaling behavior of GATK and elPrep on a variety of servers with different hardware resources. This also allows us to investigate the dollar cost in relation to the expected runtime speedups when using a particular server setup.

We additionally present a performance comparison of the previous elPrep 4 release with elPrep 5 in the S2 Appendix.

#### Validation

Since elPrep 5 produces output identical to GATK 4 (meaning it produces binary identical BAM and VCF files), we do not perform non-computational performance benchmarks where we compare the sensitivity and accuracy of elPrep 5 with relations to the variants it detects. The technical details how we achieve this are discussed in the S1 Appendix.

#### Data sets

We benchmark both a whole-exome and whole-genome data set because we expect that the impact is different for both kinds of data sets. Both data sets are downloaded from public repositories. We downloaded a 30x whole-exome data set identified as NA12878, accession SRX731649, from the 1000 Genome Project [17], and the 50x NA12878 Platinum whole-genome from accession PRJEB3381 [18].

We downloaded unaligned FASTQ data that we then aligned using BWA mem [16]. For the whole-exome data, we used the hg19 reference genome, because we can then use the hg19-compatible BED file that comes with the sample. The whole-genome data was aligned to hg38. The only significant difference between the pipelines to process the whole-exome and whole-genome data is that the whole-exome pipeline uses the BED file with captured regions for targeted sequencing.

#### Server and software versions

All of our benchmarks were executed on servers available via Amazon Web Services (AWS). The scaling experiment, specifically, was run on a variety of instances listed in Table 1. The table shows the AWS name of the instance, the number of virtual CPUs, the amount of virtual RAM, and the dollar cost per hour to rent the instance (on-demand pricing). The instances are all equipped with Intel Xeon Platinum 8175M processors. All of the instances are configured to run Amazon Linux 2. For comparing the raw computational performance of elPrep to GATK, we specifically used the m5.24xlarge instance.

**Table 1.** AWS instances used in our benchmarks. Prices for EU (Frankfurt) Sep. 2020.

| Instance | vCPU | Memory (GiB) | Cost |
| --- | --- | --- | --- |
| m5.large | 2 | 8 | \$0.115/hour |
| m5.xlarge | 4 | 16 | \$0.230/hour |
| m5.2xlarge | 8 | 32 | \$0.460/hour |
| m5.4xlarge | 16 | 64 | \$0.920/hour |
| m5.8xlarge | 32 | 128 | \$1.840/hour |
| m5.12xlarge | 48 | 192 | \$2.760/hour |
| m5.16xlarge | 64 | 256 | \$3.680/hour |
| m5.24xlarge | 96 | 384 | \$5.520/hour |

In terms of software, we used elPrep 5.0.0 compiled with Go 1.15, elPrep 4.1.6 compiled with Go 1.15, gatk-4.1.3.0 using Java 1.8.0_144, and bwa-0.7.17.

## Results

We next discuss two different benchmark experiments to predict and measure the impact on computational performance when executing variant calling pipelines with elPrep 5.

### Performance comparison GATK 4 and elPrep 5

Our first experiment consists of benchmarks that compare elPrep 5 and GATK 4 directly in terms of runtime and resource use. We show results for both a whole-exome and a whole-genome sample. All benchmarks were executed on the m5.24xlarge server available for rent through Amazon Web Services, which has 96 vCPUs and 384 GiB RAM. It is a server optimized for parallel execution, which we want to evaluate both in GATK and elPrep.

#### Whole-exome results

The benchmarks for our whole-exome data set are shown in Fig 1. The figure shows three graphs from left to right comparing runtime, peak RAM use, and disk use for both GATK 4 and elPrep 5. For GATK, the graphs show the measurements for the different steps in the pipeline. In case of the runtime and disk use, the measurements are stacked to sum up the total. In contrast, there are only individual measurements for elPrep, which merges the execution of all five steps.

**Fig 1.**
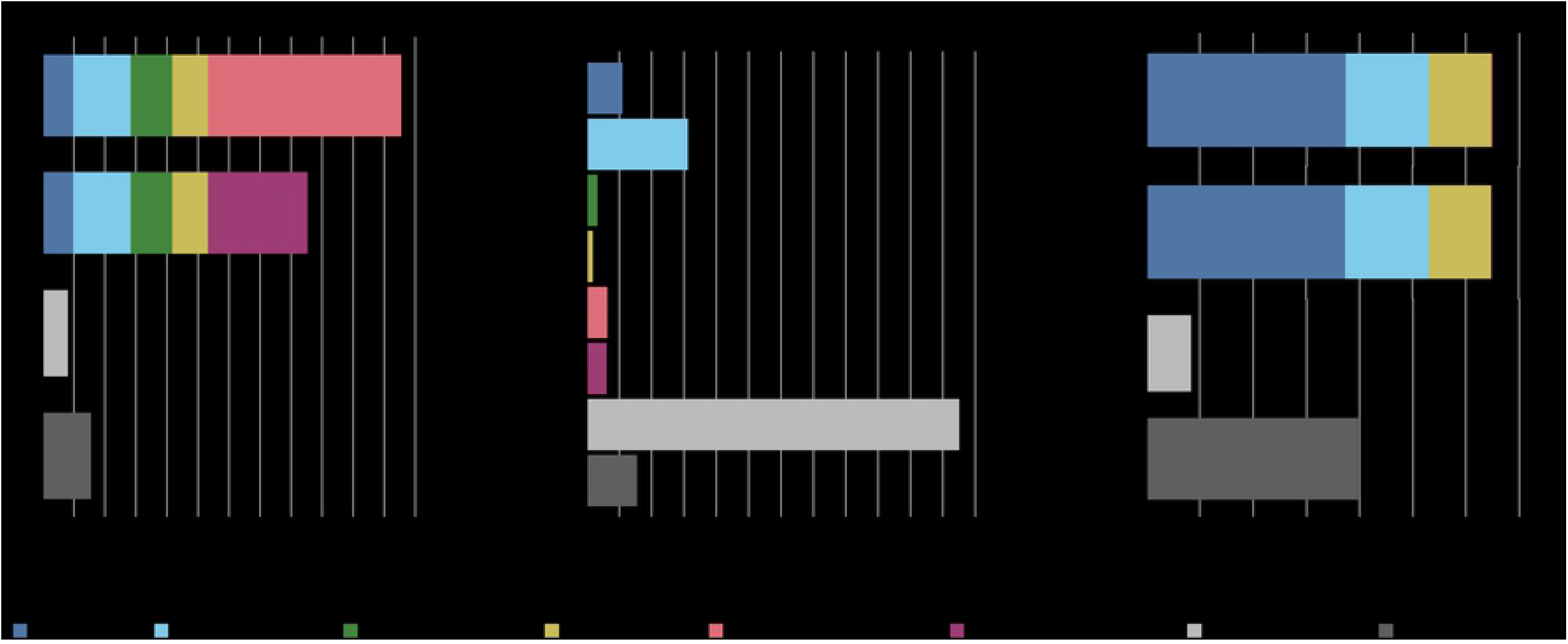
WES benchmarks. Runtime, peak RAM, and disk use in GATK 4 (Java/Intel mode) vs. elPrep 5 (filter/sfm mode). The elPrep filter mode is 11 to 15x faster than the GATK modes, using ±4x more RAM, and less than 15% of the disk space GATK uses. The elPrep sfm mode is 6 to 7.5x faster than GATK 4, uses roughly half as much RAM, and uses less than half of the disk space GATK uses. The output of elPrep 5, i.e., BAM, VCF, and metrics, is identical to the output of GATK.

The graphs show results for two execution modes for both GATK and elPrep. In case of GATK, we show the results for both the standard haplotype caller algorithm (tagged “Java”) and the parallelized version (tagged “Intel”) that relies on OpenMP multithreading and Intel AVX vector instructions for speeding up the runtime. As we previously explained, the results of the two modes differ, with some users preferring the Java mode over the Intel mode for reproducibility reasons.

For elPrep, we benchmark two execution modes, namely the filter and sfm modes. The elPrep filter mode attempts loading all input data into RAM to avoid intermediate I/O to disk, whereas the sfm mode splits up the data by chromosomal regions, yet operates without information loss [1, 2]. The filter mode is useful for processing smaller data sets such as whole exomes or shallow whole genomes more quickly than the sfm mode, but uses more RAM. Both elPrep modes produce outputs (BAM, VCF, and metrics) that are identical to the outputs produced by the GATK Java mode.

The total runtime of the GATK runs is the sum of the runtimes for the individual pipeline steps. The elPrep filter mode executes the pipeline 11 to 15 times faster than the GATK Intel and Java modes respectively. The sfm mode is 6 to 7.5 times faster than GATK. The peak RAM use in case of the GATK runs is determined by the MarkDuplicates step that uses most RAM. The elPrep sfm mode uses only half as much RAM as both GATK runs. The elPrep filter mode, in contrast, uses almost four times as much RAM as GATK. In terms of disk use, the elPrep filter mode performs less than 15% of the disk writes that the GATK runs perform. This is because the elPrep filter mode operates on data in memory and no intermediate BAM files are written as is the case for the GATK runs where each pipeline step is a separate GATK tool invocation. The elPrep sfm mode uses about half the disk space GATK uses. It uses more disk than the filter mode because it splits up the input data in different chromosomal regions on disk that it processes one by one to reduce RAM use.

In conclusion, both the elPrep filter and sfm modes execute the pipeline faster than GATK. The sfm mode uses both less RAM and disk space, whereas the filter mode uses substantially more RAM, but it is worthwhile considering for its runtime speed up. In fact, in the scaling experiment for whole-exome data that we discuss in the next section, we show that in the context of a cloud setup, the filter mode is overall the cheapest and most efficient mode to process the data, because, even though it uses more RAM than the sfm mode, the runtime is reduced so much that it reduces the server rental cost.

#### Whole-genome results

Fig 2 shows the results for our whole-genome benchmark. It shows graphs comparing runtime, RAM use and disk use for GATK 4 (Java/Intel modes) and elPrep 5 (sfm mode only). elPrep 5 executes the pipeline 16 times faster than the GATK Java run, and 8.5 times faster than the GATK Intel run that is parallelized. Concretely, elPrep executes the pipeline in less than six hours, whereas GATK requires more than 92 hours for the Java mode and more than 48 hours for the Intel mode. The elPrep sfm mode also uses less RAM, only 70% of the RAM GATK uses, namely 200 GB versus 288 GB in GATK. elPrep writes roughly 475 GB of data to disk, whereas the GATK runs write roughly 675 GB to disk, so elPrep also reduces disk use to roughly 70% of what GATK uses. The BAM, metrics, and VCF outputs of elPrep 5 are identical to the outputs from the GATK Java run.

**Fig 2.**
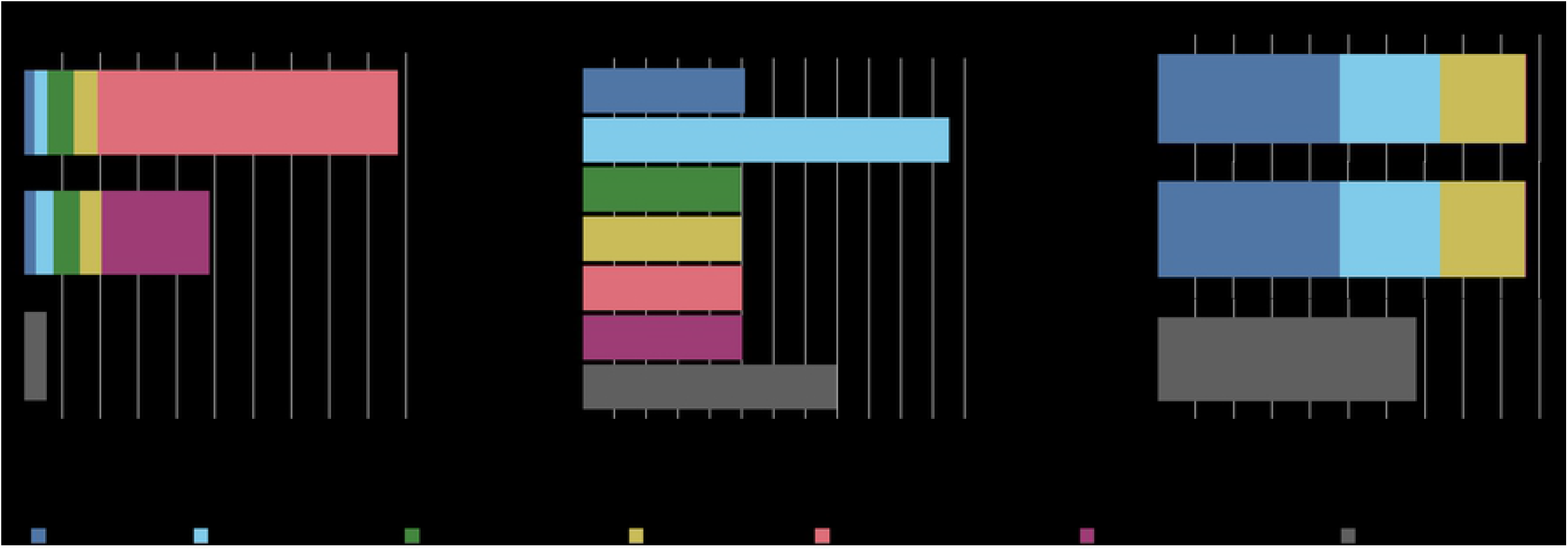
WGS benchmarks. Runtime, peak RAM, and disk use in GATK 4 (Java/Intel mode) vs. elPrep 5 (sfm mode). elPrep executes the pipeline 8.5 to 16x faster than GATK 4, depending on the haplotype caller mode used in GATK. elPrep uses only 70% of the peak RAM and disk use that GATK uses. All elPrep outputs (BAM, VCF, and metrics) are identical to the output GATK (Java) produces.

### Scaling benchmarks on Amazon Web Services

Our second experiment consists of a scaling experiment on Amazon Web Services (AWS) where we execute the pipeline on different servers with varying CPU and RAM resources, as shown in Table 1. Concretely, the servers range from having 2 to 96 vCPUs and 8 to 384 GiB RAM. Of course servers with more resources are more expensive to rent, resulting in renting costs per hour that range from $0.115 to $5.5.

The goal of the scaling experiment is to show how elPrep 5 scales compared to GATK 4, i.e., how well does each software package perform in terms of reducing runtime when it is offered access to more hardware resources? We additionally calculate the rental costs for each of the runs because we want to show what speedup can be expected for what cost. Ideally, if software scales well, the higher cost of renting servers with more computational resources can be compensated by the reduction in runtime needed.

#### Whole-exome results

Fig 3 shows the scaling experiment on AWS for the whole-exome sample. On the left it shows the runtime graph for GATK (Java/Intel) and elPrep (filter/sfm). The runtimes are shown on different AWS servers ordered from left to right by increasing CPU and RAM resources. On the right, the figure shows the cost calculated per run.

**Fig 3.**
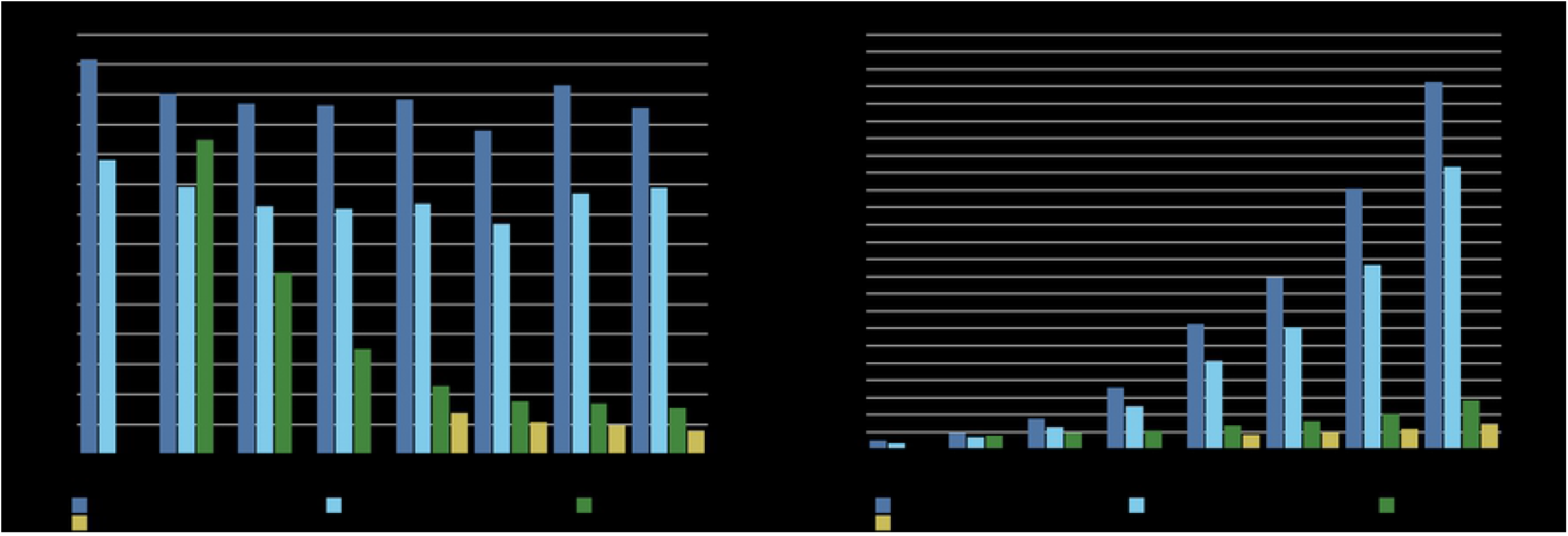
AWS WES benchmarks. The runtime and cost for running the variant calling pipeline on a variety of AWS servers, ordered by increasing CPU/RAM resources. The GATK (Java/Intel) runs show minimal improvements in terms of runtime with more resources, whereas the elPrep (filters/sfm) runs scale nearly linearly. Because of its good scaling, the cost of using elPrep on more expensive servers with more resources is countered by the reduction in runtime.

The benchmark was run for both GATK modes for variant calling, namely the original haplotype caller algorithm (Java) and the parallel version included by GATK (Intel). For both versions, GATK scales somewhat, as runtime seems to generally improve with an increase of resources up to the m5.12xlarge instance. In contrast, both elPrep runs (filter and sfm) scale much better, with the runtime nearly halved for each increment of resources. If we relate this to the cost graph on the right, we see that for the GATK runs, the cost rapidly increases with each server increment. That is because the software does not scale well, and the additional cost of a bigger server adds up to a much more expensive run than on a weaker server. In contrast, because the elPrep software scales, the cost on a bigger server is compensated by the reduced runtime.

The AWS scaling graphs can be interpreted in different ways to think about what it costs to achieve a particular runtime speedup. For example, the fastest elPrep run, filter mode on m5.24xlarge, is roughly ten times faster and 5 times cheaper than the fastest run with GATK, Intel mode on m5.12xlarge. The cheapest run overall is with GATK, Intel mode on m5.large, but it is more than 12 times slower than the fastest elPrep run and barely 4 times cheaper. For two times the price of the cheapest GATK run, you get a run with elPrep, sfm mode on m5.2xlarge, that is two times faster. If matching the output of the original haplotype caller algorithm (Java mode) is priority^1^, then the graph shows it is actually cheaper and faster to run the job with elPrep, cf. the run on m5.8xlarge with elPrep filter mode is slightly cheaper and almost nine times faster than the GATK Java mode run on m.5xlarge. In general, it is a trade off between desired runtime, price, and reproducibility of the outputs.

#### Whole-genome results

The results for the whole-genome AWS scaling benchmark are shown in Fig 4. We again show results for both GATK execution modes (Java/Intel), but only the sfm mode for elPrep. We also reduced the number of servers on which we ran the experiments because we already know the GATK software does not scale well from the whole-exome experiments. Note that no results were included for the smallest AWS server (m5.large) as we were unable to run GATK 4 on this server. An interesting observation is that the elPrep run on m5.24xlarge is cheaper than the GATK Java run on m5.2xlarge (around $32 versus $45), because the elPrep run is almost 14 times faster, reducing the runtime from almost 80 hours to less than six hours.

**Fig 4.**
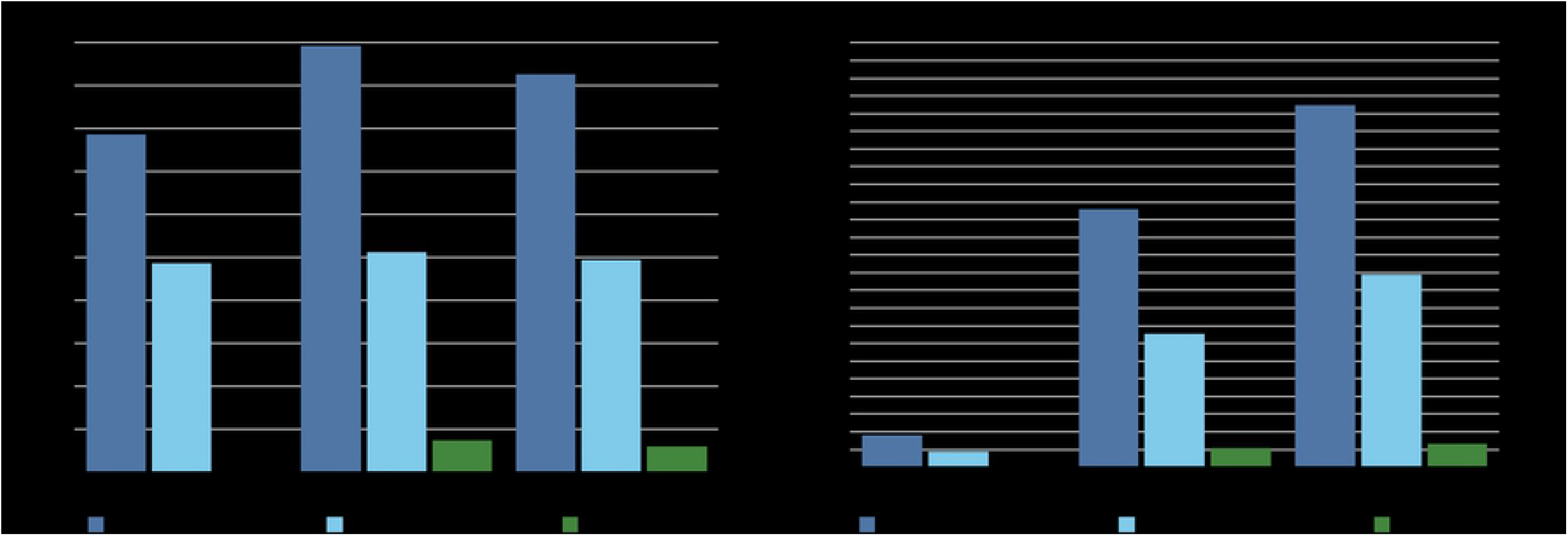
AWS WGS benchmarks. The runtime and cost for running the variant calling pipeline on a variety of servers. Similar to the WES experiment, the GATK runs (Java/Intel) show minimal scaling behavior, whereas adding additional CPUs and RAM does improve the runtime when using elPrep. The cost for GATK steeply increases because of its poor scaling whereas the cost for elPrep remains more or less stable across servers. The fastest run with elPrep is 14 times faster than the fastest run with GATK Java mode, and cheaper. The fastest elPrep run is more than 8 times faster than the fastest GATK Intel run, and only 1.4x more expensive. elPrep produces identical outputs to the original haplotype caller algorithm in the GATK Java mode.

### Additional benchmarks

We next present a number of additional benchmarks that discuss a variation of the GATK Best Practices pipeline for variant calling without base quality score recalibration.

#### Removing base quality score recalibration

The variant calling pipeline recommended by the GATK Best Practices consists of five steps, but it is not uncommon that the base quality score recalibration and application steps are removed because they are computationally demanding [21, 22], and the GATK team is currently publicly discussing that they may drop these steps from the GATK Best Practices altogether. We therefore ran the variant calling pipeline without BQSR steps to see their impact on computational performance, both with elPrep and GATK.

Fig 5 shows the results for the whole-exome sample and Fig 6 for the whole-genome sample. For the whole-exome sample, elPrep speeds up the pipeline 9x to 13x using the filter mode, whereas the sfm mode speeds up the pipeline 6.5x to 9x. For the full pipeline (Fig 1), in comparison, we see speedups around 11 to 15x and 6 to 8x for the filter and sfm modes respectively. For the whole-genome sample, elPrep (sfm only) speeds up the pipeline 5.5x to 14x compared to GATK, whereas for the full pipeline (Fig 2) we have speedups between 8.5x and 16x. This confirms our previous observation that the benefit of using elPrep increases as the number of steps in a pipeline increases [1], as elPrep’s strength lies in its execution engine that merges and parallelizes the execution of multiple steps.

**Fig 5.**
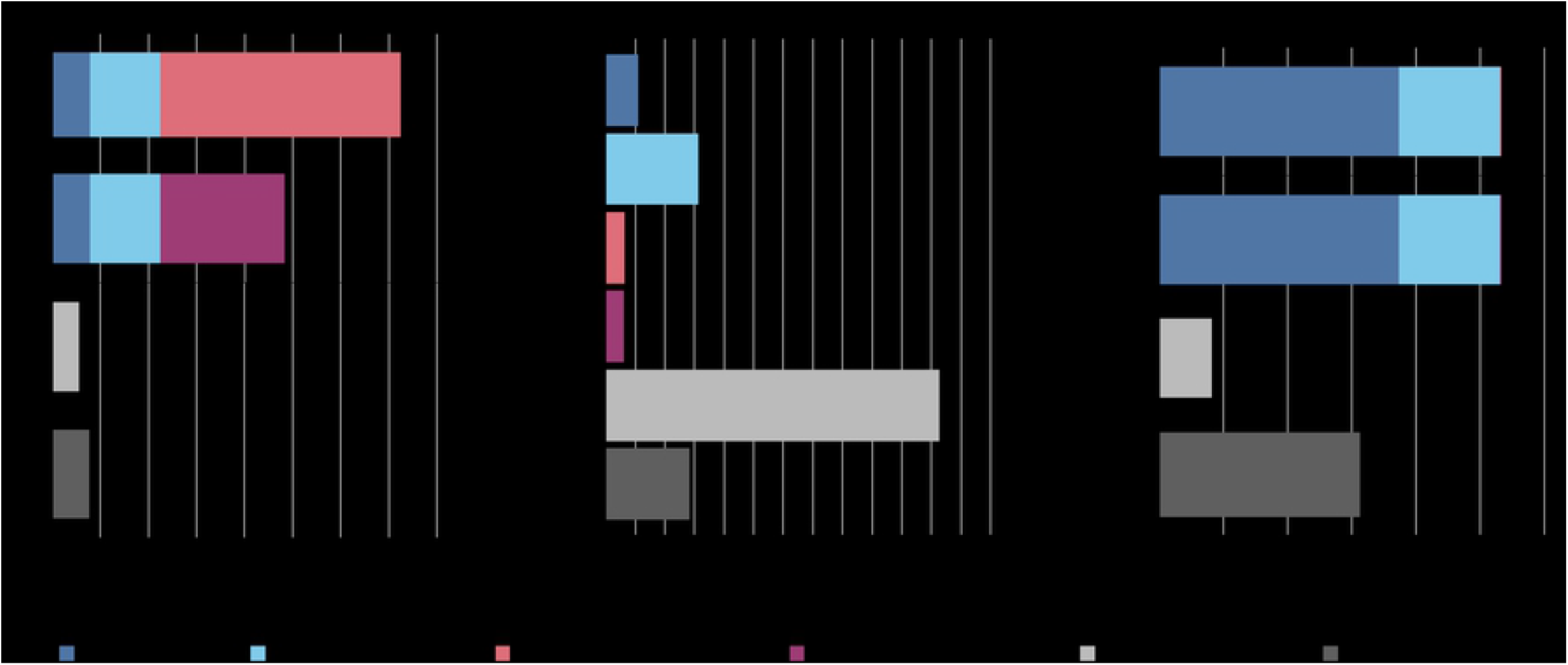
WES benchmarks without BQSR. Runtime, RAM, and disk use in GATK 4 (Java/Intel mode) vs. elPrep 5 (filter/sfm mode) on a pipeline that consists of sorting, duplicate marking, and variant calling. The elPrep filter mode is 9x to 13x faster than the GATK Intel and Java modes respectively, using ±3.5x more RAM and ±15% of the disk space GATK writes. The elPrep sfm mode is between 6.5x and 9x faster than the GATK modes, using ±90% of the RAM GATK needs, and ±60% of the disk space. The elPrep outputs are identical to the GATK Java mode outputs.

**Fig 6.**
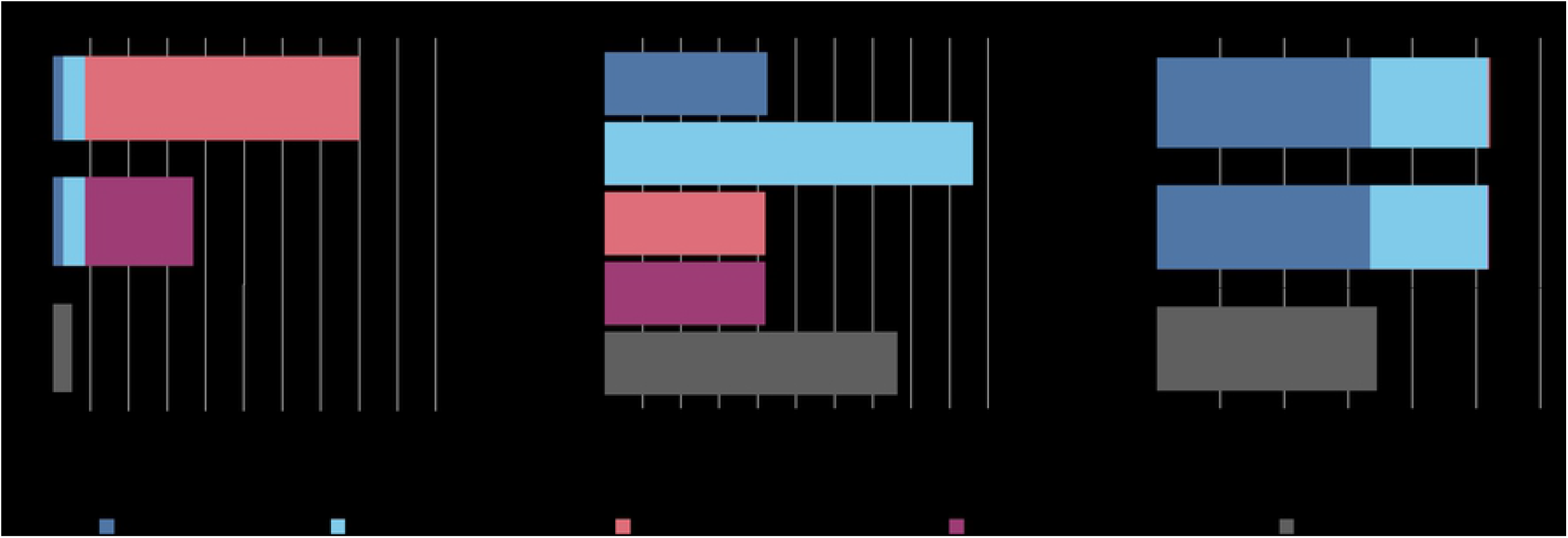
WGS benchmarks without BQSR. Runtime, RAM, and disk use comparing GATK 4 (Java/Intel mode) vs. elPrep 5 on a 3-step variant calling pipeline. The elPrep run is 5.5x to 14x faster than the GATK runs, uses ±80% of the RAM GATK uses and ±70% of the disk space. The elPrep outputs (BAM, VCF, metrics) are identical to the GATK Java mode outputs.

Another interesting way to interpret the graphs is to think about the additional cost of adding BQSR in terms of runtime and resource use. For the whole-genome sample, if you use GATK to execute the full variant calling pipeline, it is between 1.16x and 1.34x slower (Java/Intel mode) versus executing the pipeline without BQSR. In contrast, with elPrep, executing the full pipeline with the elPrep filter mode is 1.15x slower than running without BQSR. Concretely, with elPrep this is a difference between ±5 and ±6 hours of total runtime, whereas for GATK, removing BQSR reduces the runtime by ±12.5 hours for the whole-genome benchmark. In terms of RAM use, there is not much difference between the GATK runs as duplicate marking is the dominant step for memory use. We see a similar result for the elPrep runs with and without BQSR. For disk use, the GATK run with BQSR uses 1.3x more space than the GATK run without BQSR. Similarly, elPrep uses 1.4x more disk space when executing BQSR compared to running the pipeline without.

The results for the whole-exome sample are similar. For the GATK runs, running with BQSR is around 1.3x to 1.4x slower than running without. For elPrep, running with BQSR is between 1.15x (filter mode) and 1.6x slower (sfm mode). The RAM use is similar for all runs, because duplicate marking is the dominant factor here as well. For the disk use, GATK runs with BQSR need around 1.2x more space, and elPrep runs need around 1.1x more space for the sfm mode, and ±70% of disk space in the filter mode that avoids disk I/O by loading all data in RAM.

## Discussion

We can make a number of observations based on the benchmark results we reported in the sections above. elPrep 5 speeds up the execution of variant calling pipelines both for whole-exome and whole-genome data sets. The expected speedup depends on the elPrep execution mode used, which can be 1) *filter mode*, which favors keeping all data in RAM during execution, or 2) *sfm mode*, which operates on the data split by chromosomal regions on disk to limit peak RAM use. Using filter mode, elPrep speeds up the execution for whole-exome data by 11x to 15x, while sfm mode speeds it up by a factor 6.0x to 7.5x. For whole-genome data, which we only tested using sfm mode, we see between 8.5x and 16x speedup.

In general, elPrep uses both less RAM and disk resources than GATK, except when using the filter mode, which uses around 4x more RAM than GATK. This is by design, as the filter mode maximizes avoiding file I/O by keeping data into RAM to achieve optimal speedup. In the AWS scaling experiment, we also saw that, while renting a server with more RAM is more expensive per hour, the speedups with elPrep reduce the runtime so much that it can actually be cheaper to run pipelines on a bigger server.

In comparison to the benchmarks for whole-exome data, elPrep 5 shows more speedup for whole-genome data (sfm mode). This is because variant calling, which is the new feature of elPrep 5, typically takes up a larger portion of the overall runtime of a pipeline for whole-genome data. The pipeline on whole-exome data namely takes into account the regions targeted during sequencing, limiting the variant calling to use only those regions. Hence more time is spent proportionally on variant calling for whole-genome data and there is more computation for elPrep to speed up.

The benchmarks also show that elPrep’s optimization strategy – that consists of merging the execution of multiple pipelines steps, parallelizing their execution, and avoiding file I/O – is more effective than optimizing individual steps. This is illustrated by our benchmarks of GATK that show both results for running the pipeline with the original GATK haplotype caller algorithm and a parallel version thereof that is also bundled with the GATK release. While the GATK approach to optimise the single variant calling step does reduce the overall runtime (by less than 2x), elPrep shows a much better overall speedup (up to 16x). This confirms the effectiveness of elPrep’s architecture, verifying again our claims made in earlier work [1, 2, 7].

Using elPrep in general pays off more when executing pipelines with multiple steps. Our benchmarks of the variant calling pipeline without BQSR steps illustrate this. The runtime for the latter pipeline is reduced less compared to speeding up a pipeline that includes all steps (up to 14x instead of 16x). On the other hand, adding BQSR adds far less actual runtime in elPrep than in GATK. For example, a full variant calling pipeline with elPrep needs about 6 hours for a 50x WGS, whereas dropping BQSR reduces the runtime to a bit more than 5 hours. In contrast, with GATK, BQSR adds more than 12 hours to the runtime. The argument for dropping BQSR because it is computationally expensive [21,22] may therefore be worth reconsidering when using elPrep.

Our scaling experiments on AWS, where we ran both GATK and elPrep on a variety of servers with different CPU and RAM resources, show that elPrep scales much better than GATK. While GATK benefits from some parallelization across individual pipeline steps, it only shows limited runtime reduction when given more hardware resources, whereas the cost rapidly increases. In contrast, elPrep scales nearly linearly, so that running on a more expensive server with more resources almost halves the runtime, stabilizing the cost. These results show in general that speeding up software through parallelization does not necessarily have to be more expensive provided the software scales well, which elPrep does.

An important aspect that we do not evaluate with our benchmarks, is the cost of waiting for results when using a tool that is slower than elPrep. For example, elPrep reduces the runtime of a 50x WGS pipeline from 48 to 96 hours to less than 6 hours. We think that is a considerable reduction in real time. We have seen that the speedups elPrep achieves allow our users in practice to improve on their turnaround time for their research or medical tasks that rely on the analysis of sequencing data. Perhaps this hidden cost reduction is at least as important as what our benchmarks can measure.

## Conclusion

elPrep is a software package for fast analysis of sequencing data. It is a drop-in replacement for tools such as SAMtools, Picard, and GATK for overlapping functionality, as it produces identical BAM, VCF, and metric outputs. The new version, elPrep 5, introduces variant calling based on the haplotype caller algorithm from GATK. This makes it for the first time possible to completely execute the GATK Best Practices pipeline for variant calling [4, 5] with elPrep. Our benchmarks show elPrep 5 speeds up the execution by a factor 8 to 16x for this pipeline compared to GATK 4. Concretely, elPrep 5 executes the variant calling pipeline in less than 6 hours for a 50x coverage whole-genome sample, and needs less than 8 minutes for a 30x whole-exome sample. elPrep achieves these speedups using algorithmic innovations and parallelization, runs on regular Intel Xeon servers without specialized accelerators, and uses less RAM and disk resources than GATK. It is released using dual licensing, as an open-source project, but with the option of a premium license with support. For more information, we would like to point the reader to our extensive documentation online, previous publications [1, 2, 7], or published use cases [8–15] in addition to this paper.

## Supporting information

**S1 Appendix. HaplotypeCaller in elPrep 5.** We describe in detail how we reformulated the haplotype caller algorithm from GATK 4 as a parallel algorithm that fits into the elPrep 5 framework, as well as the numerous challenges we faced for guaranteeing identical output to GATK 4.

**S2 Appendix. Benchmarks comparing elPrep 4 and elPrep 5.** We discuss benchmarks that compare elPrep 4 and elPrep 5 in terms of runtime, peak memory use, and disk use, when executing a pipeline that consists of four steps to prepare data for variant calling.

## Acknowledgments

This research received funding from the Flemish Government (AI Research Program). This research also received funding from Flanders Innovation & Entrepreneurship (VLAIO) via the ATHENA project.

## Footnotes

1 As previously explained, the GATK Intel mode output differs from the GATK Java mode, with some users preferring the original mode. elPrep produces results identical to the GATK Java mode.

